# A metacommunity of complex microbial communities shows high spatial synchrony, ecological memory, gradual succession and invasion resistance

**DOI:** 10.64898/2026.11.17.767327

**Authors:** Ruyu Gao, Gorkhmaz Abbaszade, Shuang Li, Florian Schattenberg, Christine Süring, Kathrin Stückrath, Erik Schönfelder, Volker Grimm, Susann Müller

**Author notes:** shared first authorship Corresponding author: Susann Müller, Department of Applied Microbial Ecology, Helmholtz Centre for Environmental Research - UFZ, 04318 Leipzig, Permoserstr. 15, Germany.

## Abstract

Complex microbial communities are increasingly applied in biotechnological and health-related environments, yet maintaining their diversity and relative community composition over extended periods remains challenging. While dispersal can counteract the divergence of spatially separated communities, it remains unclear whether controlled mass transfer can sustain compositional synchrony in highly diverse communities over extended periods, whether this synchrony persist after dispersal is interrupted, and how robust such communities are to invasion.

Here, we investigate these questions using a loop-design reactor (LDR) comprising five local wastewater microbial communities connected by controlled mass transfer. The experiment covered multiple rounds of alternating mass transfer and insular phases over a time frame of 318 days, different mass transfer intensities, and the introduction of defined microbial invaders. Community dynamics were monitored at a high temporal resolution by single-cell flow cytometry and complemented by metagenomic analyses of selected samples to characterize taxonomic compositions and functional potential.

A Bray-Curtis-based Synchrony Index revealed that controlled mass transfer established high compositional synchrony among five local communities, which persisted for more than 150 generations. Although the community structures underwent gradual succession, the local communities remained synchronized, demonstrating parallel rather than divergent temporal trajectories. Interruption of dispersal initially promote divergence, whereas during later insular phases high compositional similarity persisted in the absence of mass transfer, consistent with ecological memory. Introduced microbial species failed to establish within resident communities, indicating high invasion resistance. Community functional potential remained largely stable throughout the experiment, although phosphorus accumulation failed during phases of high mass transfer.

Together, these findings demonstrate that controlled mass transfer can maintain long-term spatial synchrony of complex microbial communities while allowing gradual community succession. Repeated connectivity promoted compositional similarity beyond periods of active dispersal while maintaining invasion resistance and functional stability, highlighting controlled dispersal as a promising community-level strategy for managing complex microbial communities.

**Highlights:**

- Metacommunity theory was experimentally validated in a long-term complex and connected microbial ecosystem.
- Mass transfer-driven dispersal reduced stochasticity and promoted spatial synchrony among local communities.
- Synchronized communities resisted invasion while undergoing slow compositional succession.

## Introduction

Assembling and maintaining the proportions of cell types in complex microbial communities is demanding, as differences in generation times, resource requirements and competitive abilities can lead to shifts in composition over time. Under unchanging environmental conditions, particularly when community members compete for the same limiting resource, competitive interactions may progressively reduce diversity and may result in the dominance of a superior strain, consistent with the competitive exclusion principle (Gause, 1934).

Therefore, the application of microbial communities in biotechnological or health-related environments needs to address two major challenges. First, strong selective pressure may reduce community diversity and, under certain conditions, lead to the dominance of only a few or even a single strain. Second, even when community diversity can be maintained to some extent, initially similar communities may diverge in structure when maintained in isolated environments, despite being maintained under identical conditions. Such divergence can arise from stochastic ecological processes, including drift, extinction, as well as small difference in local environments (Liu et al. 2019).

There are several approaches to control the composition and stability of microbial communities. In low-complex communities consisting of only a few strains, community assembly can be designed based on defined interactions among individual members. Time, frequency, and both competitive and cooperative interactions have been exploited to promote community stability (Li et al. 2023, Yang et al. 2023; Coyte et al. 2015).

In high-complex communities, however, such interaction-based control becomes considerably more challenging because numerous competitive, cooperative, and other ecological interactions occur simultaneously. Targeted manipulation of individual species interactions is limited and control strategies are required that operate independent of detailed knowledge of community members and their interactions (Zhou et al. 2017, Burz et al. et al. 2023, Čaušević et al. 2024, Jiminez et al. 2026).

Among other proposed microbiome engineering solutions (Albright et al. 2022) one promising approach for controlling complex microbial communities is the manipulation of dispersal among spatially separated local communities, as occurs, for example, in connected wastewater (WW) treatment systems (Vucic et al. 2021), or structured host-associated microbiomes (Sorbara et al. 2022, Zou et al. 2020). The ecological framework underlying such spatially connected communities is provided by the metacommunity theory (Leibold et al. 2014). A metacommunity consists of multiple local communities that are linked by dispersal of organisms to different degrees. The strength of dispersal interplays with deterministic and stochastic processes, which can facilitate the formation of comparable local communities. We previously applied metacommunity theory to develop the loop-design reactor (LDR), in which five local microbial communities were connected by controlled mass transfer. In contrast to stand-alone-reactors, spatial connectivity within the LDR enables stabilization of a natural community over approximately 40 d (Li et al. 2022). Multiple diversity metrics demonstrated increased synchronization among the five local communities while maintaining community diversity. Measures of ecological stability, including constancy and resistance, together with the maintenance of community function, further supported the stability effect of the LDR (Baselga 2010, Liu et al. 2020).

A prerequisite for investigating the dynamics of complex microbial communities is the ability to monitor changes in community structure at sufficient high temporal and spatial resolution. Microbial flow cytometry provides such an approach by generating single-cell fingerprints of microbial communities (Koch et al. 2013, Haange et al. 2020) based on physiological properties of individual cells (Müller 2007, Cichocki et al. 2020). The fingerprints enable rapid characterization of community structure without requiring the identification of individual taxa both on the local and regional scale. Complementary, metagenomics analyses were performed on a selected subset of samples. Though metagenomics did not provide the same temporal resolution as high-frequency flow cytometry, information on taxonomic composition and the functional potential of the microbial communities are provided.

In this study, we extended the operation of the LDR to investigate the long-term stability of spatially connected complex microbial communities. We specifically aimed to determine (i) whether compositional synchrony and α-diversity can be maintained over extended periods despite continuous biomass washout, and whether synchronous patterns of community succession emerge; (ii) whether the introduction of selected microbial species disrupts synchrony, diversity and stability of the established local communities; and iii) which taxa and functional potentials characterize the communities and whether community assembly and the emergence of dominant taxa are governed primarily by deterministic or stochastic processes, particularly when dispersal among local communities is interrupted.

## 2 Materials and Methods

### 2.1 Origin and cultivation of the wastewater microbial community

A natural wastewater (WW) sample was collected from a full-scale wastewater treatment plant (WWTP) in Eilenburg, Germany (51°27’39.4’’N, 12°36’17.5’’E). After batch cultivation in a mixture of 98% synthetic wastewater and 2% peptone for 24 h at 30°C, 10 mL of the preculture were inoculated into five parallel, continuously operated bioreactors within a looped mass-transfer design (Li et al., 2022). Detailed information on the microbial community cultivation process is provided in Supplemental Information S1.

### 2.2 Experimental design

The loop mass-transfer design (LDR) serves to synchronize and stabilizes microbial communities over extended time ranges (Li et al. 2022). The design comprised five local reactors (L1-L5) that were interconnected with a loop to a sixth reactor, designated as the regional pool (R). A schematic representation and the operation of the LDR is detailed in S2.1 and S2.2 and illustrated in Supplementary Figures SF2.1 AB.

The microbial community synchronizing experiment was divided into three distinct rounds, with a cumulative duration of 318 d. Round 1 was conducted over a period of 110 d, and comprised the following phases: insular phase (IS) A (no recycling, starting on day 0), recycling phase (RC)10% (starting on day 26), RC50% (starting on day 47) and IS B (no recycling, starting on day 89). The data from Round 1 were obtained from Li et al. (2022) and were used for comparison with follow-up experiments in Rounds 2 and 3, which were performed in this study. The experiments in Rounds 2 and 3 were conducted over a period of 208 d. Round 2 encompassed the IS I phase (commencing on day 0) and the RC50% phases (I, II and III, commencing on days 24, 59, and 108, respectively). Round 3 began with IS II on day 123 and proceeded with RC10% on day 158, RC50% on day 168 and IS III on day 187. Within each phase, the initial seven days (which correspond to five times the volume exchange of a reactor) were designated as the adaptation period, which was followed by a balanced period during which the abiotic parameters were expected to remain relatively constant. As illustrated in Figure 1, the sequence of events unfolded across three distinct rounds. The rationale underlying the establishment of distinct RC50% phases is determined by the addition of selected species (S11 and SF11.2).

**Figure 1.**
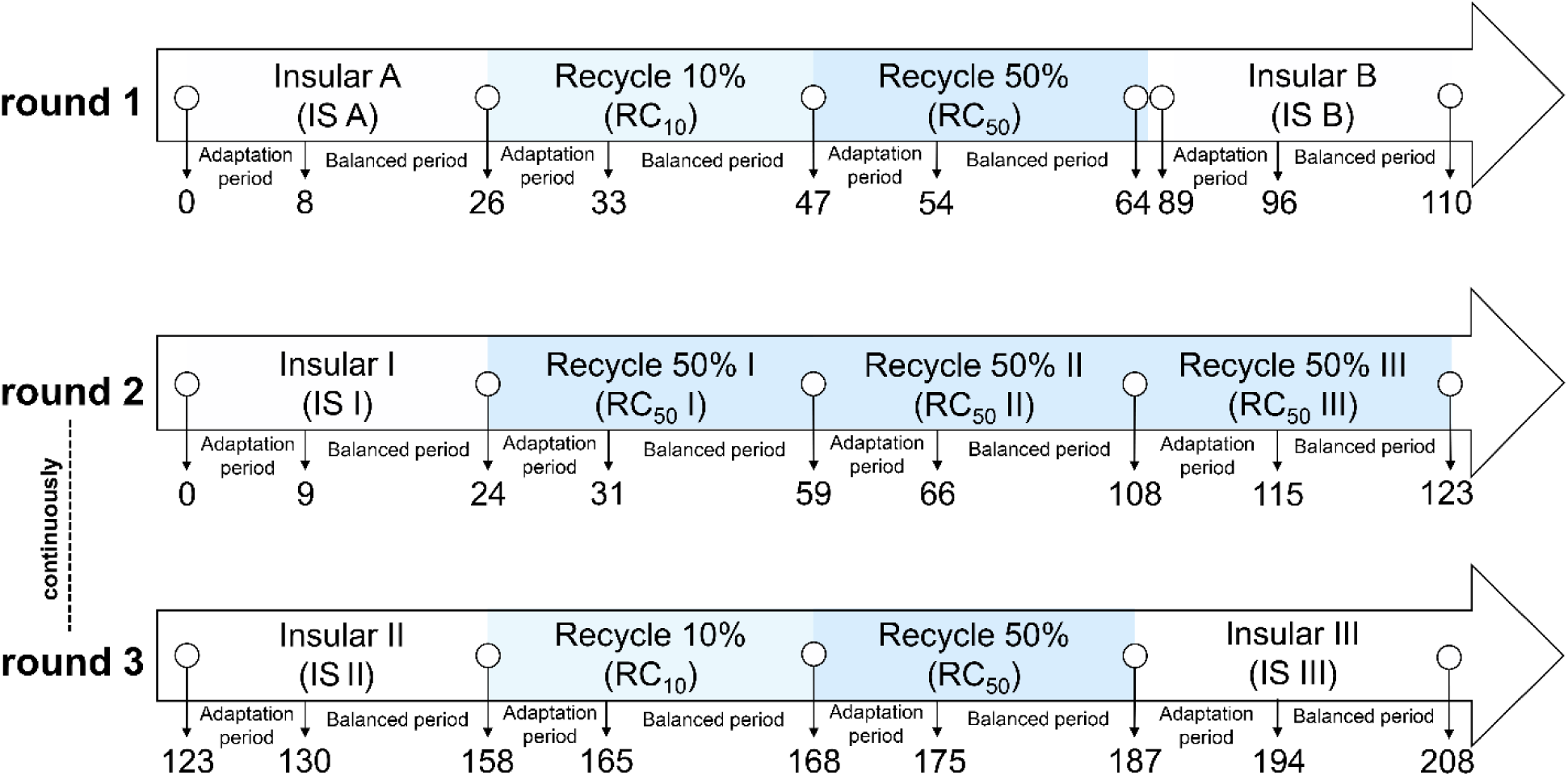
Timeline illustrating the design and sequence of the three rounds of continuous LDR cultivation. The details for the selected phases for Round 1, 2, and 3 are explained in S2.2.

### 2.3 Sampling and parameter measurement

A total of 1,226 samples were collected from the six reactors. A comprehensive analysis was conducted on each sample to ascertain the levels of abiotic and biotic parameters. The optical density at 600 nm (OD₆₀₀, path length = 5 mm) was measured using a spectrophotometer (Ultrospec 1100 Pro, Amersham Biosciences, Little Chalfont, UK). The pH and electrical conductivity (EC) were measured using a pH sensor (EL 20, Mettler-Toledo, Greifensee, Switzerland) and an EC meter (Inolab Cond7110, WTW, Germany), respectively. The dissolved oxygen content, expressed as percentage of air saturation, was monitored using an oxygen meter (OXY-1 SMA) outside of reactors, equipped with a Pst3 sensor spot (PreSens Precision Sensing GmbH, Regensburg, Germany). The detailed procedures for analyzing abiotic parameters and measured values are delineated in S2.3. The specifics of the analytical procedures for biotic data and measured values are provided below and in S3. All data were measured twice and the average value used for evaluation.

### 2.4 Cell number determination

The determination of absolute cell numbers was conducted by employing a CyFlow-Space cytometer in conjunction with the True Volumetric Absolute Counting mode (Sysmex Partec GmbH, Münster, Germany). A fresh 35 μM SYTO9 (Thermo Fisher Scientific, Eugene, OR, USA) stock solution was prepared on ice. A 50-μL stock solution was added to 950 μL of a diluted sample (final 1.75 μM), vortexed, and incubated 15 min at room temperature (RT). The measurement rate was adjusted to a range of 1,000–1,500 events s^-1^. A fluorescence cell gate was defined based on red vs. green SYTO9 fluorescence to exclude beads and noise. Absolute cell concentrations (cells mL^-1^) were obtained from events within the cell gate and corrected by the dilution factor. Data are available at Supplementary Dataset (SD) 4 (https://zenodo.org/records/24311916). Details are described in S3.3.

### 2.5 Flow cytometric analysis

Each sample from the Rounds 1-3 was processed to assess microbial community composition and dynamics by flow cytometry. A total of 365 samples were retrieved from a previously published data set (Round 1, Li et al. 2022). An additionally 861 samples were collected in Round 2 (513 samples) and Round 3 (348 samples, S3; .fcs data SD 1-3: https://zenodo.org/records/24311916).

Cells from Rounds 1-3 were fixed with paraformaldehyde, washed in phosphate buffered saline (PBS) and finally stored in 70% ethanol at −20 °C. A defined number of cells was stained with 4’,6-diamidino-2-phenylindole (DAPI) for subsequent measurement using a MoFlo Legacy Cell Sorter (Beckman Coulter, Brea, CA, USA). The procedures for cell fixation, staining, instrument operation, cytometric measurement, and corresponding control samples are outlined in S3.1, S3.2 and S3.4 (Cichocki et al. 2020; Li et. al. 2022). Cytometric data were collected as two-dimensional (2D) dot plots of DAPI fluorescence vs. FSC, providing information on DNA content per cell and cell size. The cell gate (SF3.2) was utilized to eliminate the noise and calibration beads prior to the initiation of data analysis. The cell clusters (or gates (G)) were identified using the multivariate Gaussian mixture model of FlowEMMi v2 (Bruckmann et al. 2022). A unified gate template containing 48 gates was generated based on the calculated gates and a few additionally implemented gates not found by the model. The gate template was applied to all 1,226 samples of Rounds 1-3, yielding 58,848 gates that stand for and were used to identify and track subcommunities consistently. The automatic gating procedure is explained in S3.5.

### 2.6 Evaluation of single cell data

Cells within a gate were referred to as a subcommunity (SC). Cell abundancies per SC were then used for subsequent bioinformatics analyses. In this study, dominant SCs were defined as gates exceeding 1% relative cell abundance among the 48 identified gates (S4).

The quantification of stability properties and the calculation of diversity indices (α-, intra-β, inter-β, and γ-diversity) followed the pipeline described by Li et al. (2022). These analyses were performed in RStudio (version2025.09.1+401; Posit Team, 2025) using R (version 4.5.0). A nonmetric multidimensional scaling (NMDS) based on Bray-Curtis dissimilarity of relative cell abundances per SC and correlation analyses were performed by the vegan package (version 2.7.2; Oksanen et al. 2025) and Hmisc (version 5.2.4; Harrell 2025), respectively. The ratio of turnover and nestedness for β-diversity partitioning was computed using the betapart package (version 1.6-1; Baselga & Orme 2012, Liu et al. 2020). NST analysis was estimated using the NST package (version 3.1.10; Ning et al. 2019). Data handling and visualization were conducted using reshape2 (version 1.4.5; Wickham 2007), and tidyverse (version 2.0.0; Wickham et al. 2019), including dplyr (version 1.2.0) for data wrangling. Graphical visualizations were generated using ggplot2 (version 4.0.2) together with R packages patchwork (version 1.3.2; Pedersen, 2025) and ggsignif (version 0.6.4; Ahlmann-Eltze & Pati, 2021). Heatmaps were generated using ComplexHeatmap (version 2.18.0; Gu et al. 2016), together with circlize (version 0.4.17; Gu et al. 2014). Box plots were generated using OriginPro 2026 (OriginLab Corporation, Northampton, MA, USA). Detailed descriptions of all statistical analyses are provided in S5 to S8. The R scripts are available at Zenodo: https://zenodo.org/records/24311916. Similarity percentage (SIMPER) analysis was conducted in PAST 4 tool (Hammer et al. 2001) to quantify average Bray-Curtis dissimilarities between groups and to identify the SCs contributing most strongly to observed compositional differences. For this test, SCs with relative abundances below 1% were excluded from the analysis.

### 2.7 Metagenomics analysis

A total of 56 samples were collected from the six reactors L-L5 and R, where the samples were taken from the final timepoint of each phase in Round 2 and 3 (Figure 1). Genomic DNA was extracted and its quality and quantity were evaluated. The samples were then sent to Eurofins Genomics (Germany) for sequencing by Illumina NovaSeq 6000 (S9.4.1, steps 1-3). The raw sequencing data were subjected to a series of processing steps, including adapter removal, quality assessment, trimming and removal of non-bacterial DNA. The resulting sequences were evaluated according to their taxonomic profiling and functional annotation (S9.4.1, steps 4-6). Raw metagenomic sequencing reads have been deposited in the NCBI Sequence Read Archive (SRA) under BioProject accession PRJNA1494921.

## 3 Results

An LDR was used to establish a metacommnunity, comprising five interconnected, continuously stirred bioreactors (L1-L5), which were operated under balanced conditions. The sixth reactor acted as the regional pool (R). The dynamic structural changes of the six communities were analyzed cytometrically at the single-cell level over a period of 208 d by taking 861 samples, which were evaluated alongside the samples from Li et al. (2022; 365 samples over 110 d). The ecological interpretation of the single-cell data was based on measuring 200,000 cells per sample in order to create a cytometric fingerprint. Each of them was then classified into 48 gates, which are indicative of SCs (SF 3.2). Finally, data from 58,848 SCs were available to characterize the trajectories of the respective six communities within the metacommunity over 318 d in an LDR.

### 3.1 Looped mass-transfer successfully synchronized microbial communities over extended periods of time

An overview of the time scale and operation of the LDR, with the data from Li et al. (2022, Round 1) as reference data is shown in Figure 1. Subsequent to cytometric sample handling, the resulting data underwent processing through non-metric multidimensional scaling (NMDS), an orientation method based on relative cell abundance per SC and Bray-Curtis dissimilarity analysis. The NMDS analysis encompassed the insular phases (IS), as well as the adaptation and balanced periods of the mass transfer phases (RC; Figure 2).

**Figure 2.**
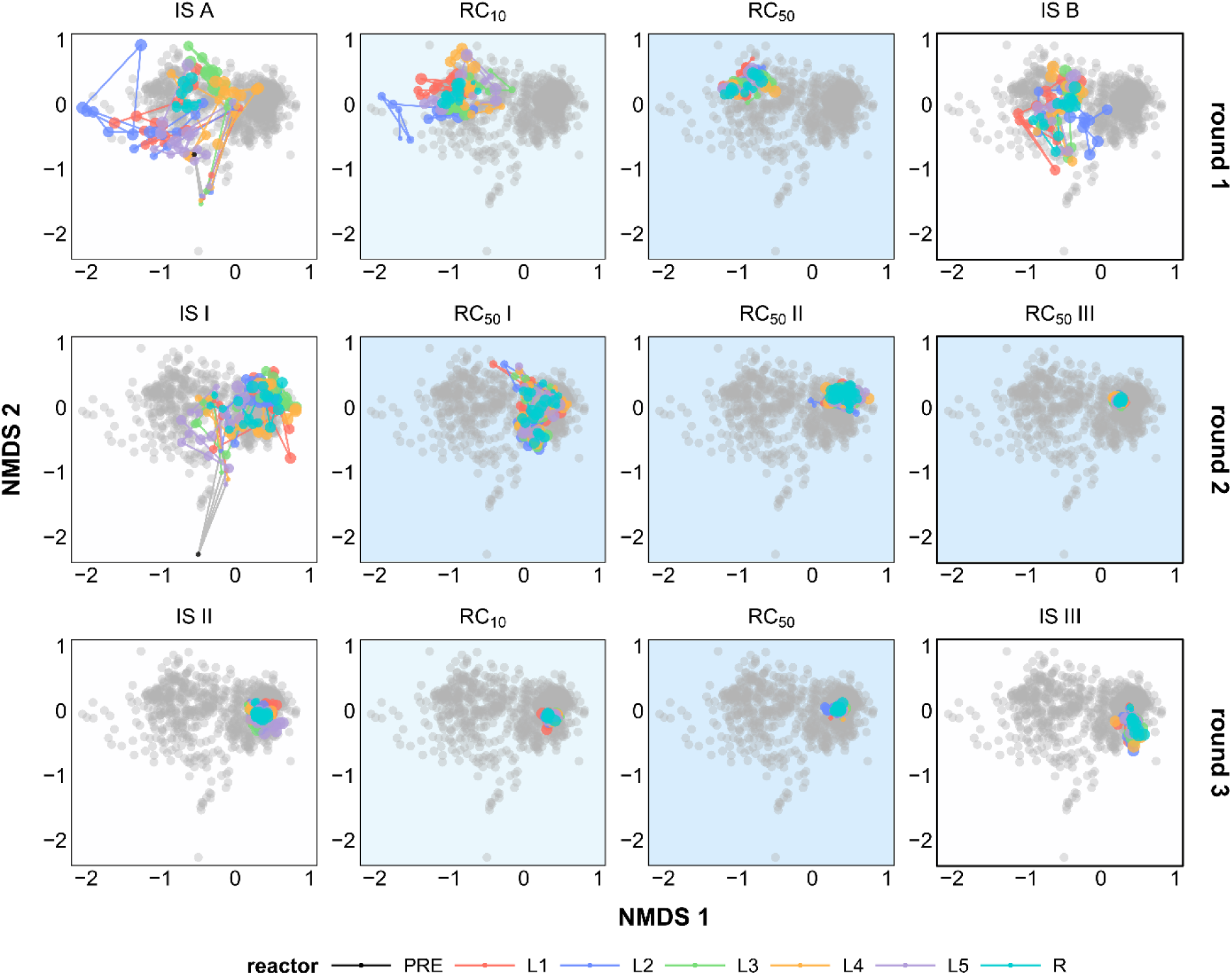
Non-Metric Multidimensional Scaling (NMDS) plots visualizing the Bray-Curtis dissimilarity between the six community structures across 1,226 samples from Rounds 1-3. Each data point corresponds a microbial community sample. The color of each point corresponds to samples from specific reactors per phase, respectively, while grey points represent samples from other Rounds and phases. Point sizes increase progressively from the beginning to the end of the respective phase (e.g., from day 1 to day 26 in the IS A phase), with successive time points connected by lines. The distances between successive points were calculated using Bray-Curtis dissimilarity. Closer points denote higher compositional similarity. Shaded backgrounds represent IS (white), RC10% (light blue), and RC50% (blue) phases.

The NMDS plot revealed significant variations in community structures among the five stand-alone reactors during the first insular phase (IS A) in Round 1. While the five reactors, operating independently, exhibited disparate trajectories, the induction of the mass transfer in the LDR was found to promote equilibrium among communities (Li et al. 2022). The equilibrium was initiated with a transfer of 10% of the biomass culture along with the medium inflow, the recycle mode RC10%, followed by a transfer of 50% of the biomass culture with the medium inflow, the recycle mode RC50%, from the regional pool (R, reactor 6) to each of the local communities (L1-L5, S2.1). A gradual decline in divergence was observed under increasing recycling rates (RC10% and RC50%) for Round 1. The structural divergence was recurring due to the subsequent disconnection of L1-L5 and R by a second insular phase IS B in Round 1.

Round 2 of the experiment proceeded with an insular phase (IS I), which once again demonstrated substantial variations between the community structures of L1-L5 and R (Figure 2). Following the activation of the LDR configuration, the subsequent recycling phases (RC50% I, RC50% II, and RC50% III) resulted in a marked decrease in community structure divergence over a period of 92 d. Round 3 of the experiment continued after Round 2 on day 123 and began with an additional insular phase (IS II), during which only minor structural alterations were observed. Subsequent implementation of the mass transfer, incorporating RC10% and RC50% phases, demonstrated a further gradual decline in variability. As anticipated, the final disconnection of the reactors in IS III resulted in an increase in the variability of the L1-L5 and R community structures, albeit to a negligible degree (Figure 2).

In order to calculate the impact of the mass transfer on community structures and their spatio-temporal equilibrium, a Synchrony Index based on Bray-Curtis dissimilarity was established (S5.6). The Synchrony Index demonstrated a marked increase alongside the number and duration of the mass transfer phases, reaching values close to 1, indicative of an extremely high degree of structural similarity among the five local communities. In Round 1, the Synchrony Index registered at 0.51 ± 0.08 across RC10%. It subsequently increased to 0.78 ± 0.05 in RC50% and rose further to 0.9 ± 0.02 in Round 2, RC50% II and III, and 0.93 ± 0.02 in Round 3, RC50% (Figure 3). Despite the disruption of the spatial synchrony among L1-L5, introduced by the insular phases, the divergence between rounds became progressively less pronounced. In Round 3, the ability of insular phases to disrupt synchrony among communities was found to be nearly nullified. The Synchrony Index during the insular phases increased from 0.3 ± 0.07 in IS A to 0.58 ± 0.1 in IS I, Round 2 and further to 0.89 ± 0.03 in IS III, Round 3. Collectively, these findings imply that recurrent mass transfer resulted in an enhancement of spatial compositional synchrony among local communities, which persisted even during subsequent periods of spatial isolation.

**Figure 3.**
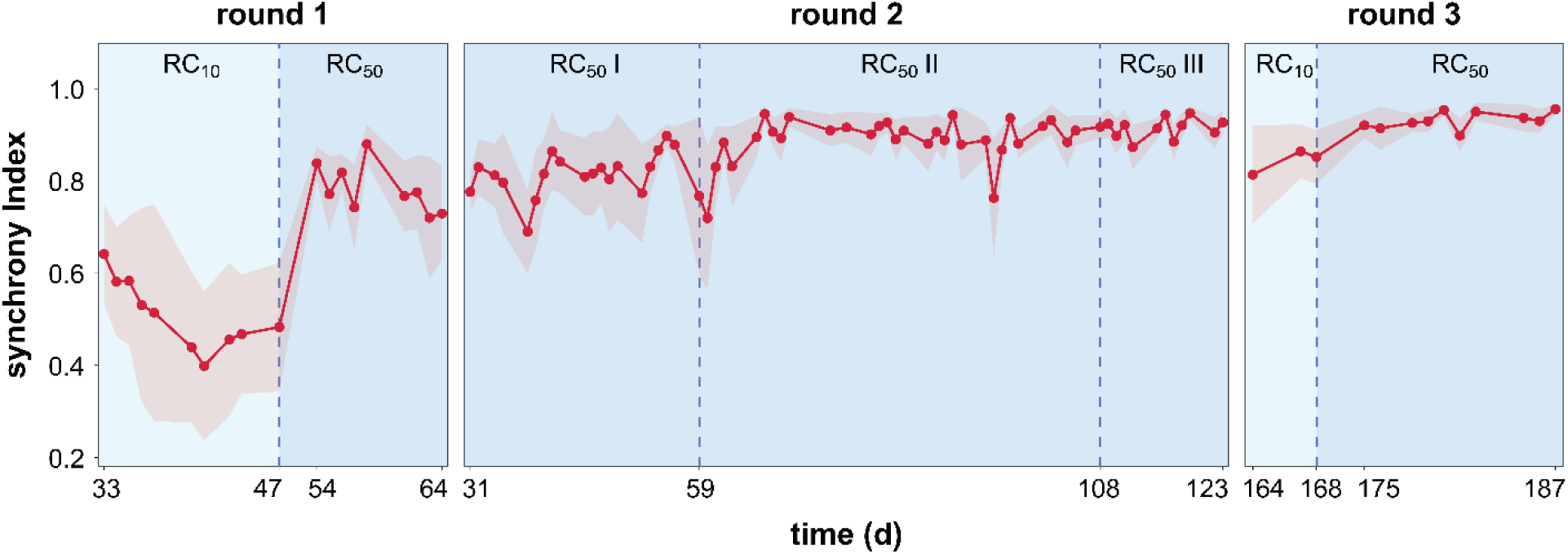
Temporal dynamics of synchrony in WW communities during the balanced periods of mass transfer phases. The red line indicates the Synchrony Index of the top 35 most abundant SCs across local communities and rounds, representing the degree of spatial synchronization by their abundance dynamics over time and at spatial scale. Shaded backgrounds represent RC10% (light blue), and RC50% phases (blue).

In order to ascertain whether the high Synchrony Index, obtained in the LDR, is exerting a detrimental effect on ecological stability, stability metrics were calculated, particularly constancy and resistance (RS) values. The constancy space is indicative for the fluctuations of the members of a community over time during the balanced periods of continuous cultivation. A low constancy value is associated with high stability and low natural fluctuations in microbiome structures. These minimal fluctuations can be attributed to the cell cycle activities of microbiome members, such as the transition between cell cycle stages due to proliferation and cell division. The constancy space was determined using a method of Liu et al. (2018). The calculated constancy space of the RC50% phase in Round 3 was selected as the reference because this was the phase at which the highest Synchrony Index was found. The cell abundance data of each SC during the balanced periods per phase was used to determine the deviation of communities in L1-L5 and R from the reference using normalized Canberra distance (S5.7). Given the presence of six communities per phase, six constancy spaces are demonstrated in Figure 4.

**Figure 4.**
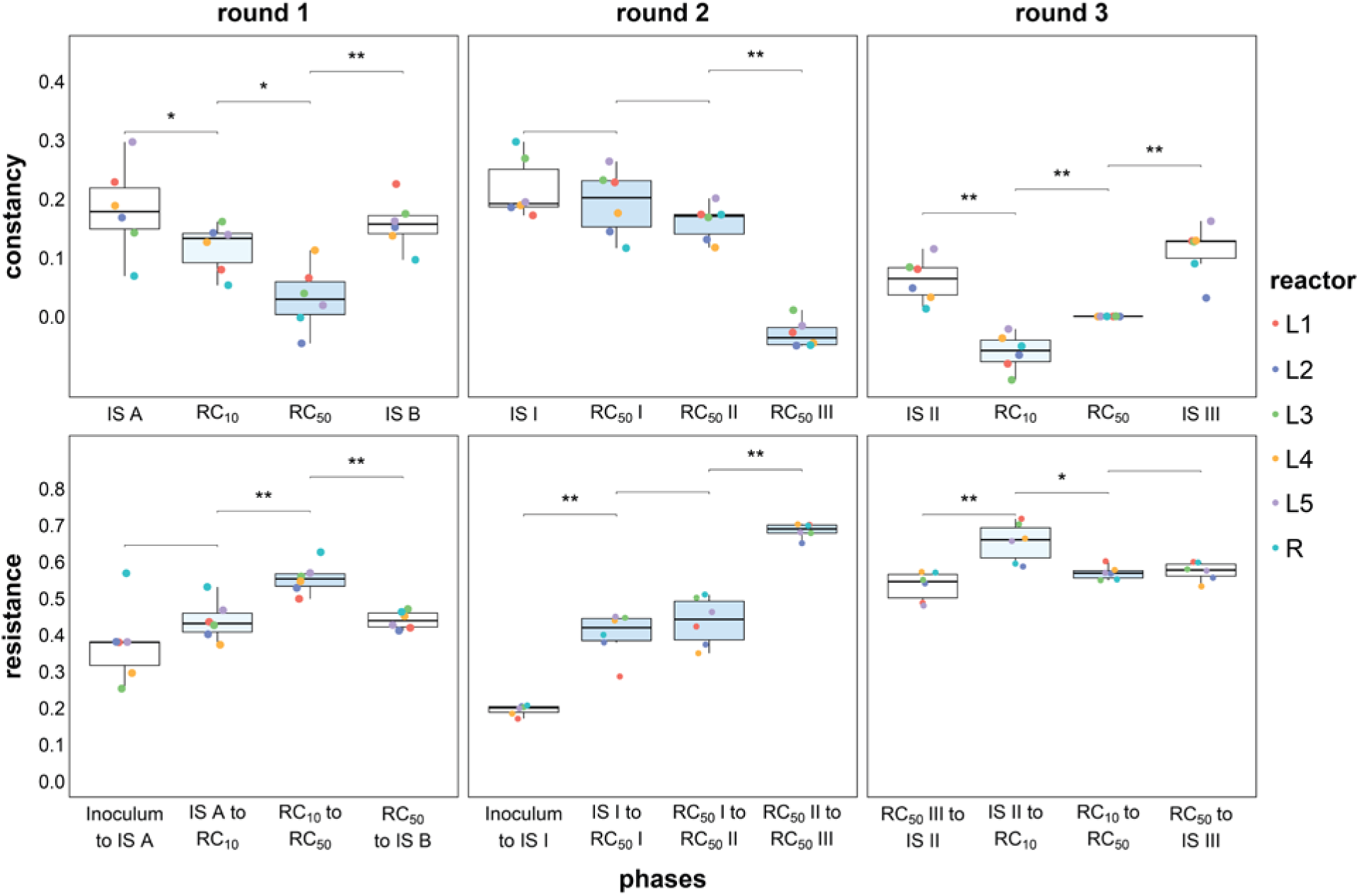
The stability properties constancy (above) and resistance (RS, below) were calculated for the balanced periods of the local microbiomes L1-L5 and the regional pool R. The box plots above illustrate the distance between aligned constancy spaces and the reference constancy space at RC50%, Round 3. Each point is representative of a constancy space per reactor community per phase. Low constancy values are indicative of unchanging community structures. The box plots below illustrate resistance values. High RS values are indicative of high resistance behavior. The stars indicate significant differences between constancy values and RS values of pairwise phases (Wilcoxon; \**p* ≤ 0.05 and \*\**p* ≤ 0.01). The shaded box plots represent the IS phases (white), RC10% phases (light blue), and RC50% phases (blue).

The RC50% III phase in Round 2 and the RC10% phase in Round 3 exhibited even more diminished values when compared to the reference constancy space, thereby supporting high stability in Round 2 and 3. Larger constancy spaces compared to the reference indicated a moderate decrease in stability, which was nevertheless minimal for all samples from mass transfer phases with a maximum constancy value of 0.19 ± 0.06 (RC50% I, Round 2). In contrast, the insular phases exhibited consistently higher constancy values compared to the mass transfer phases per Round, resulting in diminished stability. The evidence further demonstrated that over time, there was a trend of decrease in values from Round 1 to Round 3, suggesting that the microbiomes exhibited increased stability during the course of the experiment.

Resistance (RS) describes a community’s ability to stay unchanged under disturbance. In this context, the disruption of mass transfer is regarded as a disturbance, defined as a decrease or cessation of mass transfer, specifically during insular phases. The values were calculated from the balanced periods and their pairwise comparison between phases and per reactor in Rounds 1-3. RS values represent the deviation from the endpoint of a phase, e.g. IS A (day 26), to the respective endpoint of the subsequent balanced period of a phase, e.g. RC10% (day 47) (Liu et al. 2018). The highest mean RS value was found to be 0.68 ± 0.02, which was observed for the shift of RC50% II to RC 50% III, Round 2 (Figure 4). In Round 3 all RS values, including the insular phase shifts, were above 0.5, indicating a high state of resistance in communities during this round. In comparison with the mass transfer phases within each round, the insular phases consistently exhibited lower RS values and, consequently, lower resistance. Regardless, the course of the experiment revealed that the microbiomes exhibited increased RS, thereby demonstrating increased stability confirming the calculations of the constancy values (Figure 4).

In summary, the LDR repeatedly and increasingly synchronized communities, even after insular phases were implemented to disrupt synchrony. Between the balanced phases of Rounds 1-3 the Synchrony Index increased to high values. The effect was so strong that an increasing spatial synchrony was observed even between insular phases, which otherwise promote community structure variability. These findings robustly validate of the long-term feasibility of the LDR design, wherein the rescue effect strongly facilitated microbiome equalization and synchronization. Superior stability properties of constancy and resistance were evident during long-term cultivation in Rounds 2 and 3 at RC50%.

### 3.2 Microbial communities re-assemble to distinct structures among rounds

The LDR operation ensured the same levels of nutrients and carbon across phases and rounds (S2.2). Nevertheless, due to mass transfer, the cell number was increasing from the balanced phase of IS I, with a cell number (CN) of 1.63 × 10¹² cells L⁻¹ to a CN of 7.63 × 10¹² cells L⁻¹ in the balanced phase of RC50% III, which is approximately 4.7-fold (S2.3).

To demonstrate that the increases in CN do not disrupt equalization of the local communities, a SIMPER analysis was employed to compare community structure dissimilarity between balanced periods of all phases of the experiment, considering cell types (i.e., SCs) and cell abundances per SC. SCs with relative abundances below 1% were excluded from the analysis.

The inocula, coming from the precultures (S1), differed by 42.8% between Rounds 1 and 2 when the medium was flowing in (Supplementary Table (ST) 5.1). The two initial insular phases IS A and IS I established divergent structures between the communities L1-L5, exhibiting a 74.2% disparity. When comparing the dissimilarity values of the two mass transfer phases in Round 1, the disparity was 49.7%. In Round 2, for phases RC50% I, II, and III, the disparity was even lower with 41.8%, even though the time span was considerably longer (ST5.1). A comparison of Rounds 1 and 2 using the SIMPER analysis revealed a 79.5% disparity in community structures, and indeed, the structures between them were entirely different (Figure 5). Instead, an increasing uniformity was found between the mass transfer phases in Rounds 2 and 3 (33.2%), a trend that was even more evident when calculated for Round 3 alone (21.2%). Most interestingly, the structure of the communities in the two insular phases of Round 3 differed by only 33.6%, thereby implying a formation of ecological memory. All SIMPER analysis values between communities in balanced phases of Rounds 1-3 are summarized in ST5.1.

**Figure 5.**
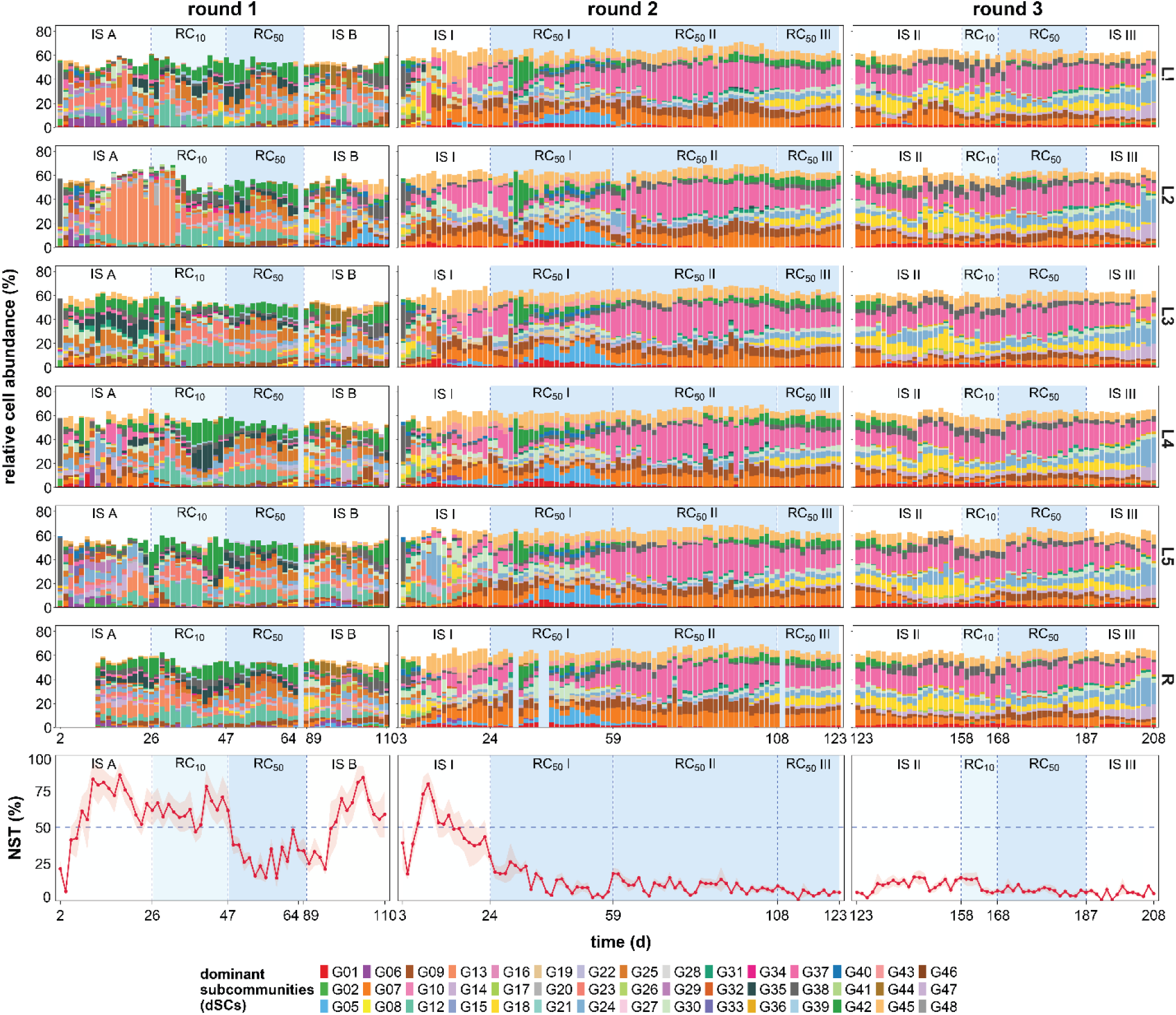
Distribution of dominant SCs and the degree of stochasticity during the process of community assembly. Above: Stacked plots of relative cell abundance of dominant SCs across the Rounds 1-3. Each column represents a different round, and the rows 1 to 6 correspond to the local communities L1-L5 and the regional pool R, respectively. Any SC that was dominant once was included in the stacked plot. Dominant SCs are indicated by distinct colors, with gates ordered numerically from bottom to top. Any gate information <1% in all samples and at all time points was disregarded. Below: The relative importance of stochastic processes in community assembly over time by comparing cell abundance changes of dominant SCs in reactors L1-L5 and R using the Normalized Stochasticity Ratio (NST). High stochasticity is represented by a high NST index. Shaded backgrounds represent the IS (white), the RC10% (light blue), and RC50% phases (blue).

The diversity indices support the data from the SIMPER analysis and were used to describe the SC establishment during the community assembly processes for all reactors L1-L5 and R. All diversity metrics were influenced by the LDR operation with exception of α-diversity. The Hill number ^0^D (Hill, 1973; Jost, 2006) reflects community richness and was in this study calculated by counting the number of the dominant SCs (S4, FS5.1). Round 1 showed a slightly higher richness (mean 14.2 ± 0.52) compared to Round 3 (mean 12.21 ± 0.21). The Hill number ^1^D encompasses evenness distributions and revealed also minimal variations between Rounds 1-3 (mean 10.49 ± 0.75 to mean 8.85 ± 0.15, FS5.2). Larger variations were found for the inter-community β-diversity measures, which captured the number of unique dominant SCs through pairwise comparison of reactor samples per day (FS5.4). The results indicated substantial structural variations among reactors L1-5 (mean 12.87 ± 2.25) and L1-5 and R (mean 11.30 ± 2.25) in IS A in Round 1. In the mass transfer phases of Round 3 the inter-community β-diversity values were low with mean values of only 1.00 ± 0.38 (L1-L5) and 1.26 ± 0.7 (L1-L5 and R). Similarly, γ-diversity (FS5.5) declined from IS A (mean 28.09 ± 3.42) to Round 2 (mean 15.71 ± 3.70), but remained nearly unchanged for the subsequent 85 d (mean 14.26 ± 1.43) in Round 3. The γ -diversity included all unique SCs between all reactors per day. Although richness remained relatively stable, the other diversity indices indicated that particular types of microbial community subgroups underwent selection during the assembly process, especially in Rounds 1 and at the start of Round 2, which resulted in a gradual convergence that sustained for over 150 generations. Notably, in Round 3, there were no differences in diversity characteristics between the insular and mass transfer phases, which supports the presence of an ecological memory effect under long-term LDR operation.

### 3.3 Stochastic and deterministic processes both shape microbial community assembly

The LDR facilitated strong dispersal through mass transfer, an operation that reinforces deterministic forces. Particular SCs such as G09, G38, and G45 were supported among L1-L5 and R (S4). Conversely, emergence of SC G18 in RC50% II in Round 2, for example, indicates the presence of stochastic forces within the LDR, which was originally designed to prevent changes to community structures. In order to ascertain the relative contributions of niche versus neutral paradigms to the structure of microbiomes during insular and mass-transfer phases, additional tools were utilized to investigate their respective community-shaping forces. The evaluation procedures encompass the dominant SCs of reactors L1-L5 and R for Rounds 1-3.

The degree of stochasticity was determined by employing the approach developed by Ning et al. (2019). The normalized stochasticity ratio (NST) has been developed as a metric to quantify the relative importance of stochastic versus deterministic processes in community assembly. Specifically, NST employs a null model representing stochastic assembly, which is compared to the assembly of the observed microbial community structure. The comparison was done by Jaccard metrics based on relative cell abundance per gate.

The NST analysis revealed that stochastic processes were particularly prevalent during the insular and subsequent phases (Figure 5, below). As the duration of the active mass transfer phases without disruption increased, the NST values decreased, reaching approximately one-fifteenth of the mean value in IS A (mean 72.8% ± 1.87%). The lowest values were observed in phases RC50% II and III in Round 2 (mean 9.29% ± 1.43%, mean 4.56% ± 1.32%, respectively) and in phase RC50% in Round 3 (mean 5.37% ± 0.76%). The NST analysis clearly corroborated the earlier finding that mass transfer reduces stochasticity (Li et al. 2022). However, the NST index also revealed that short-term mass transfer situations still harbor a high probability of being stochastic. For instance, this occurs under RC10% in Round 1, with an NST mean value of 61.60% ± 2.32% and under RC50% I in Round 2, with 10.57% ± 1.31%. It is noteworthy that the mean NST value in IS III (4.68% ± 1.54%) was analogous to the RC50% value in Round 3, suggesting again an ecological memory effect of long mass transfer phases that extended into the insular phase. These findings imply that neutral events are low during mass transfer but can exert a lasting influence on community structures, albeit one that is often imperceptible but persistent.

The findings were corroborated by intra-community β-diversity values, which are indicative of drift events in microbial communities and were counted for L1-L5 and R (S5.2, ST6.1). Drift events mirror random changes in community structures and have been shown to lead to divergent assemblies (Liu et al. 2019). A comparative analysis revealed that all insular phases exhibited elevated drift rates in comparison to the corresponding mass transfer phases. Notably, the occurrence of drift events diminished between rounds, exhibiting a decline from an average of 8.17 in RC10% in Round 1 to zero in RC10% and R50% in Round 3 (ST6.1). Stochasticity reemerged to a minor extent in IS III Round 3 (ST6.1), concurrent with the cessation of mass transfer.

While NST and intra-community β-diversity values revealed the extent of neutral behavior, particularly in the insular phases, rank-order abundance curves (FS7.1) and the Sørensen Partitioning Index (FS5.6) provided insight into the degree of deterministic forces, especially in Rounds 2-3. The rank-order abundance curves exhibited equal, S-shaped distributions of SCs with equivalent cell numbers during the IS I insular phase (FS7.1, above), while steeper slopes were observed for RC50% in Round 3 (FS7.1, below; ST7.1), indicating the predominance of deterministic forces in Round 3. In support of this assessment, the Sørensen Partitioning Index indicated a substantial decrease in turnover values with time (S5.5, FS5.6). The findings provide substantial support for a deterministic interpretation of mass transfer processes.

In summary, the NST calculations and intra-community β-diversity values suggested a substantial influence of stochastic processes. Conversely, the steep rank-abundance curves and low turnover values of the Sørensen Partitioning Index following repeated mass-transfer phases indicated the presence of strong deterministic forces, thereby supporting the maintenance of community structures.

### 3.4 Microbial community functions during insular and mass transfer phases

The primary functions of a WW community are the degradation of organic carbon, the removal of nitrogen, and the accumulation of phosphorus from WW. These processes result in the production of bacterial biomass that is enriched in phosphorus. In managed natural systems such as WWTPs, this biomass is subsequently removed as biosolids. An important question is whether the WW community can maintain these functions within the LDR or whether its configuration may affect the core functional performances. Specifically, it remains unclear whether the efficiency of the communities differs between stand-alone reactors, represented by the insular phases, and reactors operating under mass-transfer conditions, represented by the RC phases. Furthermore, it was of interest whether these functions remain stable throughout the 318-day operation of the LDR. As we observed shifts in the structural composition of the WW community between rounds, the question was rising of whether functional performance is maintained. It seems also crucial to identify the species responsible for carrying out these functions within the LDR.

To address these questions, interaction patterns derived from cytometric fingerprints (S8) were calculated and visualized and the efficiencies of carbon, nitrogen, and phosphorus removal quantified (S9.1 - S9.3). In addition, metagenomic analyses were used to characterize the functional potential of the WW communities and to link their composition with ecosystem functioning (S9.4).

#### 3.4.1. The interaction potential between SCs is highest in mass transfer phases

Spearman’s rank-order correlation coefficient rho was employed to evaluate the interaction strength and network pattern among SCs and bulk biotic and abiotic parameters. All cytometric fingerprinting data and all bulk data (FS2.2) were included in the analysis, and a strong correlation factor of I0.9I with a p-value >0.05 was chosen (S8). Correlations were estimated across L1-L5 and R in Rounds 1-3, and the following configurations were examined: SC vs. SC, SC vs. parameter (SC vs. Para), and parameter vs. parameter (Para vs. Para).

The highest total numbers of correlations were identified during mass transfer phases, particularly in RC50% II (547) and RC50% III (1016) in Round 2, and RC10% (886) and RC50% (524) in Round 3. Conversely, insular phases exhibited a markedly lower prevalence of correlations, such as IS I (257) and IS III (277, ST8.1). Correlation numbers were much less frequent between SCs and bulk parameters and even less frequent among the bulk parameters themselves, ranging from 1 to a maximum of 22 correlations per phase (ST8.1, SF8.1 and SF8.2). As an example, visualization of the interaction patterns between SCs and bulk parameters in L1-5 and R during Round 2 revealed a substantial increase in the number of both positive and negative edges from IS I to RC50% III, with the highest network connectivity observed in R (SF8.3). Notably, correlations involving bulk abiotic parameters were distributed across a wide range of SCs, suggesting the absence of robust associations between specific SCs and particular abiotic parameters, indicating that functional redundancy is common within the WW communities.

#### 3.4.2. Mass transfer phases maintain carbon and nitrogen removal but reduce phosphorus removal

A total of 1,226 samples were collected from L1-L5 and R to determine the concentrations of COD (chemical oxygen demand of the supernatant), NH₄-N (ammonium nitrogen in the supernatant), and PO₄-P (orthophosphate in the supernatant) (methods described in Section S2.3). The data for Rounds 1-3 is collectively presented in FS2.2 and DS3 (Round1), and DS4 (Rounds 2 and 3).

The CODs values were used to assess the degradation of soluble organic carbon in the WW. During this process, organic carbon is utilized as a source of energy and carbon for the synthesis of bacterial biomass (S9.1). During the insular phases, the CODs values remained close to zero and increased only slightly during the mass transfer phases. The average CODs removal efficiency was slightly higher in Rounds 2 and 3 than in Round 1, reaching 81.88% ± 9.33% (ST9.1).

The NH_4_-N values were used as an indicator of amino acid deamination during protein degradation by measuring ammonium accumulation in the medium (S9.2). During microbial metabolism, nitrogen bound in proteins and amino acids is either assimilated into bacterial biomass or released as ammonium through deamination. Under aerobic conditions, this ammonium can be oxidized to nitrite and nitrate by nitrification, thereby ensuring low levels of NH_4_-N values in the early insular phases. In contrast, during mass transfer phases, bacterial cell number (CN) increased by approximately 4.7-fold (S2.3), likely reducing oxygen availability and thereby inhibiting nitrification, which resulted in ammonium accumulation in the medium from approximately 30 mg L^-1^ to 60 mg L^-1^. Based on the total nitrogen concentration (S9.2), the NH₄–N removal efficiencies were 66.97% ± 4.58% in Round 1. For Rounds 2 and 3, the efficiencies were calculated separately for the two operational phases, yielding 79.74 ± 25.8% for the insular phases but still 77.58% ± 15.69% for the mass transfer phases (ST9.1).

The PO_4_-P values were used to assess orthophosphate uptake from the medium, reflecting phosphorus assimilation into bacterial biomass or intracellular phosphorus storage (S9.3). Aerobic conditions generally favor PO_4_-P uptake, whereas oxygen-limited conditions may reduce PO_4_-P uptake and, in communities containing polyphosphate-accumulating organisms (PAOs), promote PO_4_-P release into the medium. Accordingly, PO₄-P concentrations remained low in insular phases. In contrast, during high biomass phases, when the bacterial CN increased substantially and oxygen availability was reduced, PO_4_-P values increased to maximum 159 mg L⁻¹ (FS2.2, SD4). Therefore, the PO_4_-P removal efficiency was evaluated separately for low biomass phases, in which oxic conditions prevailed, and high biomass phases, such as the mass transfer phases. Based on the total phosphorus concentration (S9.3), the PO_4_-P removal efficiencies were 11.87 % ± 2.92 % in Round 1. For Rounds 2 and 3, the efficiencies were 77.63% ± 30.21% during the low biomass phases and 21.90 % ± 5.67% during the high biomass phases (ST9.1).

In summary, across Rounds 1-3, the WW communities in L1-L5 and R maintained consistent functional performance with respect to carbon and nitrogen removal during both the insular and mass transfer phases, whereas phosphorus removal efficiency remained limited during the mass transfer phases.

#### 3.4.3 Metagenomic functional profiles corroborate cytometric correlations and bulk measurements

Metagenomic functional profiles align with cytometric correlations and bulk measurements. From the final sampling time point of each phase, a total of 56 samples from reactors L1–L5 and R were selected for analysis (S9.4.1). Overall functional analysis revealed high pathways completeness for carbon cycling, including carbohydrate metabolism, carbon degradation, and amino acid metabolism. Pathways associated with nitrogen metabolism, particularly those involved in nitrification and denitrification, also exhibited high completeness. In addition, pathways involved in phosphorus metabolism were partly apparent (S9.4.2). The identified metabolic pathways are consistent with the functional potential expected for a WW community (FS9.1).

Functional gene profiles (S9.1), obtained from the metagenomics analysis of the 56 samples, were visualized by principal coordinate analysis for overall pairwise dissimilarity (Figure 6B). All insular phases (IS I-IS III) in Rounds 2-3 clustered closely, indicating relatively small differences in functional potential across phases. Likewise, the functional profile at the end of RC50% I (Round 2) clustered with the IS phases. By contrast, the profiles at the end of RC50% II and RC50% III (Round 2) formed distinct clusters, which but accounted for just 5.3% of the functional differences from the other cluster. This functional difference coincided with the highest number of strong correlations in the cytometric datasets (e.g., RC50% III: 780 SC-SC, 214 SC-Para, and 22 Para-Para; total: 1016; ST8.1, SF8.1). Finally, the functional gene profile proved resilient, returning to its initial functional capacity during IS II in Round 3, verified both by the metagenomic and cytometric datasets (Figure 6B, SF8.1).

**Figure 6.**
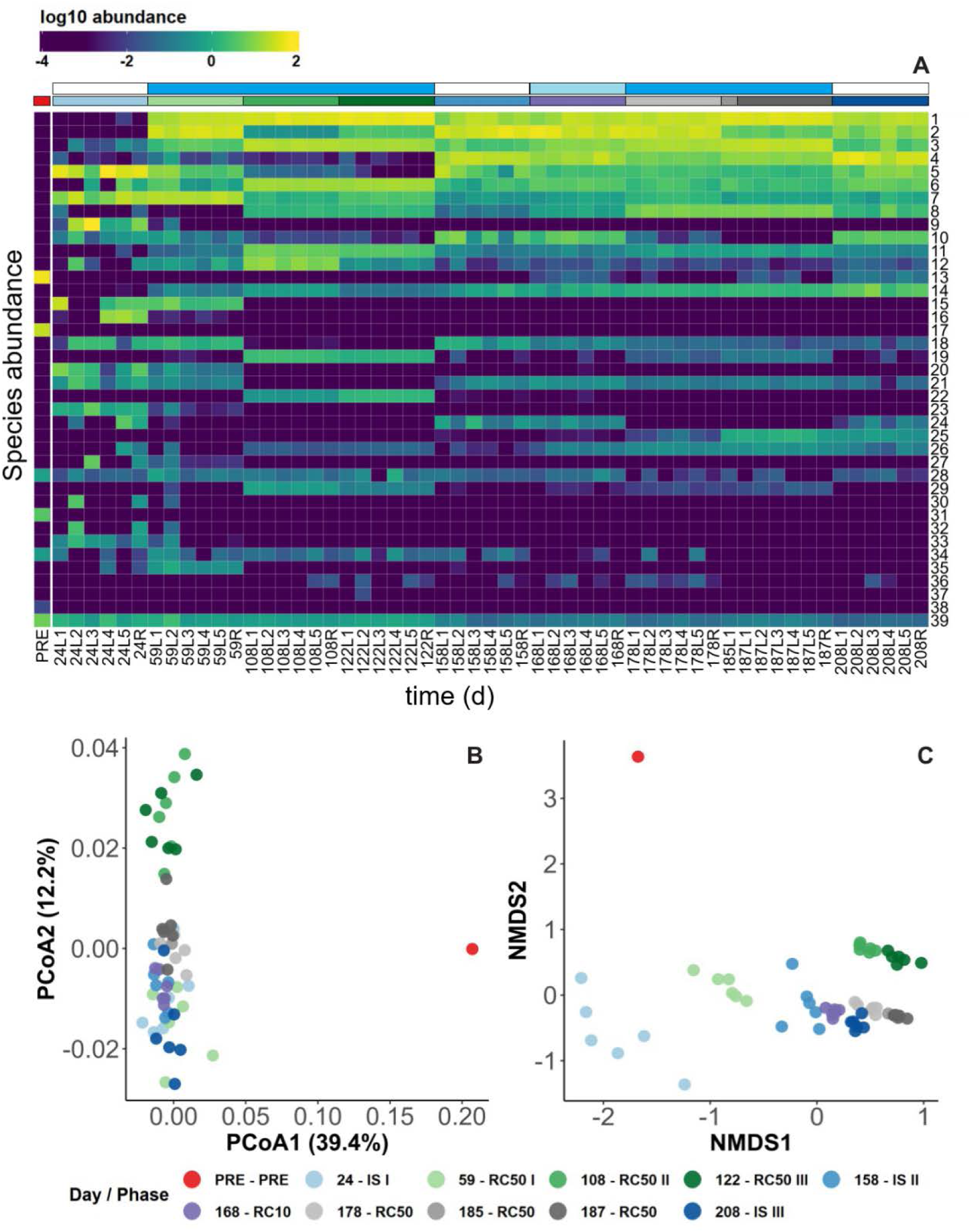
Metagenomic analysis of functional and taxonomic dynamics of the WW communities in L1-L5 and R in Rounds 2-3. A: Relative abundance data of the 35 most dominant species (species names in S10). PRE represents the preculture. Species 36-38 represent the augmented strains *P. citronellolis* P3B5 (36), *S. melonis* Fr1 (37), *E. coli* DH5α (38), and others (39). Nr. 39 includes all other remaining species combined (from 0.03% to a maximum of 6.4%). The upper left color scale indicates the log_10_-transformed relative species abundance. B: PCoA of functional gene profiles, using Bray-Curtis distance measure. The samples are grouped based on similarity of functional gene profiles. C: Non-metric multidimensional scaling (NMDS) plot of species-level taxonomic profiles, using Bray-Curtis distance measure. The samples are grouped based on the similarity in community composition. For B and C: Each point corresponds to a sample, and each color indicates samples from reactors L1-L5 and R, operated in parallel. Corresponding days and phases are shown in the upper horizontal color legend above A. The second shaded upper horizontal color legend represents the IS (white), RC10% (light blue), and RC50% phases (blue).

### 3.5 Metagenome-derived community structure reflects functional profiles and process parameters

The metagenomic data were used to determine the taxonomy of the dominant species that emerged in the LDR and dominated the WW communities for more than 150 generations. The dynamics and relative abundances of the 35 most dominant species are shown in Figure 6A. Together, these species accounted for 93.6–99.97% of the total community relative abundance, thereby representing nearly the entire community. *Aquamicrobium lusatiense* (1), *Aquamicrobium defluvii* (2), and *Reyranella massiliensis* (3) were consistently among the most abundant species throughout Rounds 2 and 3. Other persistent but less abundant taxa included *Acidovorax* sp. MR-S7 (4), *Achromobacter xylosoxidans* (5), *Flavobacterium lindanitolerans* (6), *Brevundimonas diminuta* (7), and *Parapusillimonas granuli* (8). The dominant species belonged mainly to heterotrophic classes Alphaproteobacteria and Betaproteobacteria, which are commonly detected in WW communities (S10). These taxa are generally associated with organic carbon utilization and biomass production. Classical nitrifying microorganisms were not detected among the 35 most abundant species. However, several species associated with nitrogen transformation were identified, including *Acidovorax* sp. MR-S7 (4), *P. granuli* (8), and *Stenotrophomonas nitritireducens* (10). Canonical phosphorus-accumulating organisms (PAOs) were also not among the 35 most abundant species identified.

In addition, the taxonomic profiles from the metagenomic data analysis were used to assess dynamics in WW community composition during Rounds 2-3 (Figure 6C). In contrast to functional PCoA analysis, NMDS ordination, based on species-level taxonomic profiles generated with MetaPhlAn, revealed clear temporal changes in community composition (Figure 6C). In line with the results of the functional PCoA analysis, samples from RC50% II and RC50% III (Round 2) formed a distinct cluster, clearly separated from samples of the other LDR phases. The main species responsible for this difference seem to be *Acidovorax* sp. MR-S7 (4) and *Achromobacter xylosoxidans* (5, Figure 6A). All samples from Round 3 clustered together both functionally and taxonomically, showing that the most dominant strains in all local communities within the LDR during this round had unchanging profiles over a 50-day timeframe (Figure 6 ABC).

### 3.6 The priority effects prevent colonization by exogeneous taxa

The LDR induced exceptional synchronization and stabilization pressure on the structure and function of the local communities L1-L5, despite the presence of some intrinsic stochastic forces. To further investigate the influence of external biological factors on community resistance, three exogeneous taxa (*Pseudomonas citronellolis* P3B5 (36), *Sphingomonas melonis* Fr1 (37), and *Escherichia coli* DH5α (38)) were introduced each at a maximum relative abundance of 3.7% of the WW community (Figure 6A, S11 step 4). Each strain was chromosomally-labelled with a distinct fluorescent protein, namely red fluorescent protein (mScarlet-I), yellow fluorescent protein (sYFP2), or green fluorescent protein (GFP), respectively (S11), enabling strain-specific abundance dynamics to be monitored by flow cytometry. Strain cultivation, multifluorescence flow cytometry settings, measurement procedures, and gating strategies are described in S11. The experimental procedures for single- and combined-strain additions are provided in Figure S11.2. Different strain combinations, addition levels, and exposure durations were tested (S11). Neither *E. coli* DH5α nor *S. melonis* Fr1 established within the community in any combination. Only *P. citronellolis* P3B5 was detected during six days of continuous addition while connected to the LDR (days 168-174), reaching a relative abundance of 0.3%. After discontinuation of the addition, its abundance declined to 0.001% by day 208.

The cytometric observations were supported by metagenomic analyses (SF11.3). Neither *E. coli* nor *S. melonis* was detected at the final sampling point, whereas *P. citronellolis* was detected only at very low abundance using the more sensitive MetaPhlAn settings (0.008%). A comparison between the cytometric and metagenomics taxonomic profiles is shown in SF11.4, which displays relative abundances estimated by cytometry (cra) and metagenomics (mra, Spearmans rho 0.667, p = 6.63E-06, ST 11.1). Throughout the experiment, the introduced taxa remained below 0.06% relative abundance after their addition to the established WW community, with *P. citronellolis* P3B5 also declining to near-undetectable abundance by the end of the experiment. The results indicate a high resistance of the local WW communities L1-L5 to colonization by incoming organisms, suggesting that, although the rescue effect may promote the persistence and establishment of newly incoming organisms, strong priority effects constrain their successful integration into the established communities of the LDR.

## 4 Discussion

### 4.1 Long-term looped mass-transfer enforced robust spatial synchrony of local communities

A high Synchrony Index of >0.9 was facilitated by the LDR, in which five local microbial communities were operated in parallel and were continuously connected to a regional pool (Figure 3). Flow cytometric analysis identified at least 11 dominant SCs that contributed substantially to the observed spatial synchrony. Figure 7 shows the cytometric SCs with high Synchrony Index values, quantifying the similarity of relative cell abundance trajectories for each individual SC across at least 3 local communities. Metagenomic analyses supported these findings by revealing 35 dominant taxa that remained at nearly unchanged relative proportions over more than 150 generations (Figure 6a). These dominant taxa accounted for approximately 95% of the total community abundance of which 15 taxa exceeded 1% average relative abundance and maintained highly similar rank positions across the five local communities (Figure 6a).

**Figure 7.**
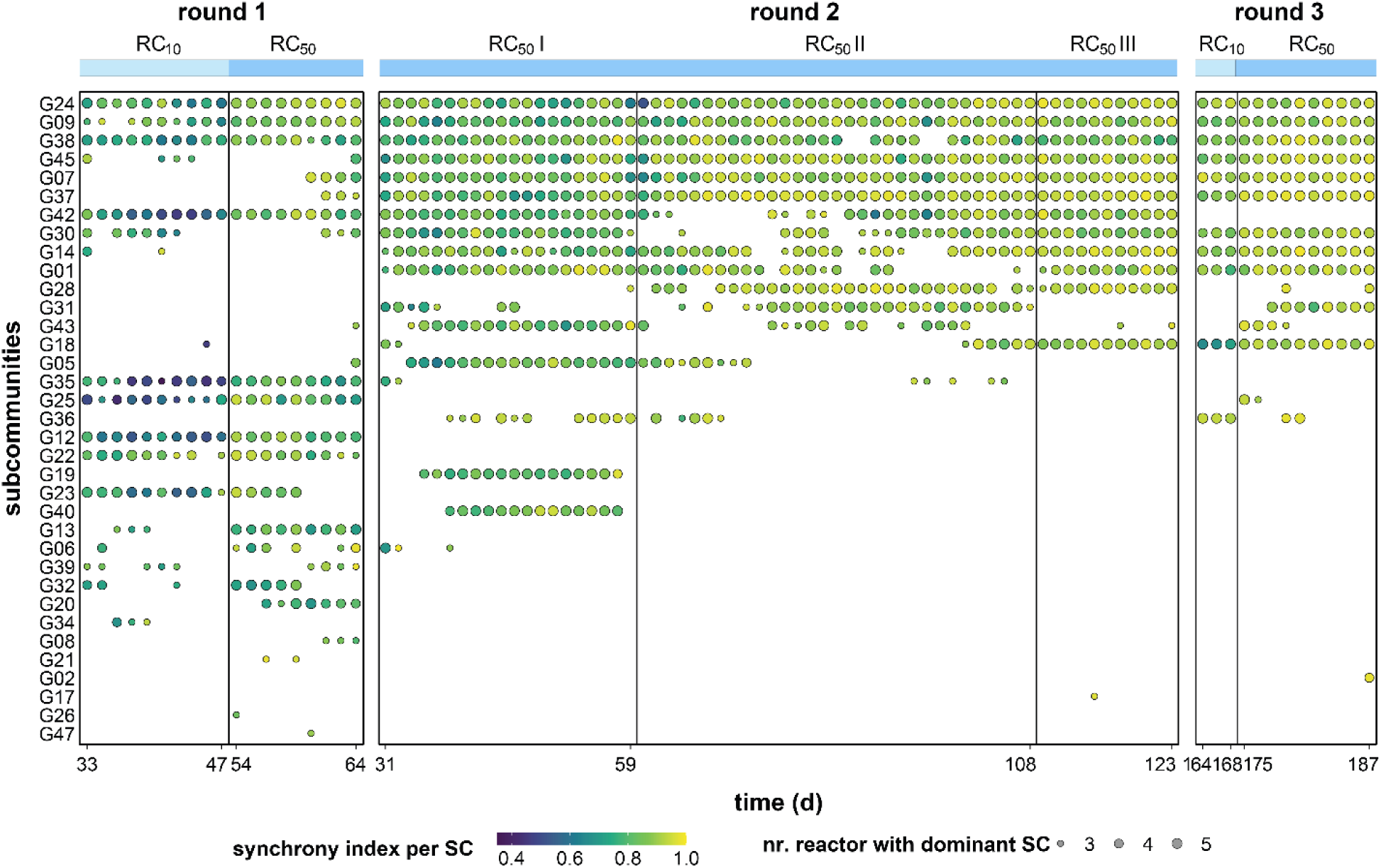
Dynamics of spatial synchrony among reactors per SC in the WW communities of L1-L5 during the balanced periods of mass transfer phases in Rounds 1-3. Only SCs with a relative cell abundance >1% are shown. Circle size indicates the number of reactors (3–5) in which a SC was dominant at each time point, while the color code represents the synchrony index of that SC across reactors. The shaded upper horizontal legend represents the RC10% (light blue) and RC50% phases (blue).

The synchronizing effect of high dispersal among local communities is well established within the metacommunity theory. Studies across macroecological systems have demonstrated in animal and plant populations that increasing dispersal among habitat patches promotes spatial synchrony, particularly when dispersal rates are high and habitat patches remain connected over extended timescales (Abbott 2011, Wang et al. 2019, Reumann et al. 2025). However, whether such synchrony can emerge and be maintained over extended timescales in complex, naturally derived microbial communities remained largely unexplored till now. Short-term synchronized microbial metacommunities have been demonstrated by Li et al. (2022), whose dataset was incorporated into the present study. Other studies have similarly focused on short-term experiments or spatial snapshot comparisons (Declerck et al. 2012; Fodelianakis et al. 2019), whereas long-term, spatially replicated experimental datasets remain scarce. Our study demonstrated that continuous high dispersal can transform spatially separated microbial communities into a single, robust, and highly synchronized metacommunity providing rare long-term experimental support for the synchronizing effect of the mass transfer framework of metacommunity theory.

### 4.2 Local communities established plateaued diversity and stability values within the metacommunity

Spatial synchrony can be caused by environmental fluctuation such as climate forces (Moran, 1953; Kendall et al. 2000) or by dispersal among local communities (Yang et al. 2022). Generally, environmentally induced spatial synchrony is expected to reduce regional stability because synchronized communities may lose their buffering capacity, especially under environmental perturbations (Larsen et al. 2025). In contrast, studies on dispersal have shown that spatial synchrony can improve stability, as observed in our system, although there are also indications that exceedingly high dispersal rates may also lead to instabilities (Abbott 2011; Yang et al. 2022).

In our study, environmental conditions were methodologically balanced, and spatial synchrony was induced by controlled dispersal. The establishment of spatial synchrony did not coincide with a decline in community diversity and stability. The individual microbial communities maintained their stability properties by the rescue effect. The growth of community members decreased due to unchanging nutrient concentrations and operational conditions, accompanied by a more than fourfold biomass increase due to mass transfer. The cytometric α-diversity values (Hill numbers ^0^D, ^1^D) converged spatio-temporally towards even values (FS5.1, FS5.2) and also the metagenomic analysis revealed no variation in dominant species abundance (Figure 6). Inter-community β-diversity values and γ-diversity values did not change with the onset of elevated mass transfer events in Round 2 (FS5.4, FS5.5). In parallel, the Sørensen Partitioning Index showed constant nestedness and low turnover values (FS5.6).

As stated in the study by Grimm et al. (1997), stability properties reflect temporal fluctuations of community members, which were determined in this study by using the two properties constancy and resistance. The two properties identified the exceptional stability of the five individual communities during mass transfer in our study. Low constancy and high resistance values were shown to be closely linked to dispersal-induced spatial synchrony also in other systems (Davidson et al. 2023). In our study, spatial synchrony was coupled with unchanging diversity and stability properties under the continuous dispersal regime established in our highly connected microbial metacommunity.

### 4.3 Long-term spatial synchrony dos not exclude ecological memory processes, gradual community succession, and metabolic function and shows invasion resistance

In metacommunity studies, reducing dispersal rates has been found to lead to strong compositional divergence (Chen et al. 2020; Yang et al. 2022). In our study, dispersal was not only reduced but the local communities were temporarily isolated for 20 to 35 generations. Under these conditions, the communities followed independent structural trajectories (Figure 2; Round 1, IS B, data from Li et al. 2022). After restarting the experiment (Round 2) with the same inoculum, in the beginning only a slightly different initial setup was observed (ISA and IS I, Figure 5, first stacked bars), which highlights the importance of deterministic environmental filtering imposed by the identical operational conditions of the LDR in shaping community assembly (Chase et al. 2011, Stegen et al. 2013, Dini-Andreotea et al. 2015). However, these differences alone cannot easily account for the fundamentally different succession of SCs observed between Rounds 1 and 2 a few days later. The different successions may partly reflect priority effects, whereby taxa that establish early influence the subsequent trajectory of community assembly (Zhou et al. 2017). Additionally, since the inoculum community originated from a complex natural WW microbiome, there might be functional redundancy, meaning that functionally equivalent organisms can easily replace each other (Li et al. 2021). Accordingly, carbon and nitrogen removal efficiencies differed only marginally between IS A and IS I (ST9.1). Nevertheless, the stochastic processes appear to have reached their full potential only after approximately 10 generations in each of the three insular phases in Rounds 1 and 2. The NST reached maximum values of 0.860 ± 0.085 (day 15, Round 1), 0.843 ± 0.082 (day 104, Round 1), and 0.800 ± 0.079 (day 8, Round 2; Figure 5). Likewise, intra-community β-diversity indicated elevated levels of ecological drift throughout the three insular phases of both rounds (FS5.3). Only the onset of the prolonged mass transfer phases in Round 2 substantially reduced stochasticity, resulting in the establishment of strong spatial synchrony across the five local communities.

However, a 35-day disruption of the long-term mass transfer phases between Rounds 2 and 3 did not result, as before between Round 1 and 2, in renewed divergence among local communities or the establishment of distinct local community structures (Figure 5). Instead, the high degree of spatial synchrony established during Round 2 persisted throughout the subsequent insular phases. The formation of intense interaction networks (FS 8.1) and the reduced influence of stochastic assembly processes (SF7.1, Figure 5) may represent mechanisms contributing to such history-depending dynamics. Thus, previous periods of connectivity appeared to influence community dynamics even after connectivity was removed, leaving a structural legacy of their previous connected state. Several ecological processes may contribute to such ecological memory (Odum, 1969, Dini-Andreotea et al. 2015, Canarini et al. 2021). For example, repeated mass transfer may progressively homogenize local species pools and thereby reduce their potential for subsequent divergence. Beyond the dominant cell types considered here, homogenization or loss of rare community members may also contribute to this effect, an aspect investigated in a parallel study (Abbaszade et al. 2027).

Importantly, this ecological memory did not imply structural constancy (Gajrani et al. 2025, Zhang et al. 2026). The spatially synchronized metacommunity continued to undergo minor but gradual compositional successions throughout Rounds 2 and 3. However, the local communities underwent these changes in parallel. Ecological memory may have therefore constrained their subsequent community development without preventing future succession itself. Such dynamics can be described as synchronous succession, in which community composition changes gradually over time while spatial synchrony among local communities is maintained.

The functional performance also remained highly synchronous among local communities (SF2.2). Consistent with the overall functional profile, metagenomic analysis also revealed no alterations in metabolic potential, except for one small one that was exclusively found during the prolonged mass transfer phases in Round 2 (Figure 6B, S9.1). Taxonomic metagenomic analysis identified species commonly associated with carbon degradation and nitrogen removal in WW, with functional annotations indicating genetic potential related to amino acid metabolism, ammonification, ammonium oxidation and denitrification (S9.4). The near absence of detectable polyphosphate-accumulating taxa within the metacommunity was unexpected (S9.4) but may reflect the low oxygen availability during later mass transfer phases (FS2.2) and agrees with the low phosphorus removal efficiencies observed during Rounds 2 and 3 (ST9.1).

To assess the invasion barriers of the spatially synchronized metacommunity, three autofluorescent bacterial reporter strains were introduced at varying proportions and time intervals (FS11.2). The inability of newly introduced taxa to establish within the already assembled metacommunity under all tested conditions suggests strong historical constraints on community assembly. Although continuous mass transfer provided repeated immigration opportunities, the established metacommunity structure prevented successful invasion of the introduced taxa, even when they were provided with a relative abundance of up to 3.7% (S11). The previous community assembly likely shaped resource availability (FS2.2) and interaction networks (S8), thereby restricting colonization by late-arriving invaders. These findings indicate that a metacommunity shaped by ecological memory exhibits strong colonization barriers, highlighting the importance of community history in assembly trajectories.

## 5. Conclusion

This study demonstrates that the LDR provides a unique experimental platform for investigating metacommunity dynamics in complex natural microbial communities. Continuous mass transfer generated a highly spatially synchronized metacommunity that maintained remarkably constant α-diversity, inter-community β-diversity, and γ-diversity over more than 150 generations when mass transfer was established. Spatial synchrony was accompanied by high community stability, reflected by consistently low constancy values, high resistance, and a marked reduction in stochasticity. Functional performance remained largely unchanged compared with the disconnected insular phases, with the exception of reduced phosphorus removal under mass transfer, which was associated with increased biomass and limited oxygen availability. Functional and taxonomic metagenomic analyses supported these observations, revealing only minor changes in metabolic potential during prolonged mass transfer and a persistent dominance of the same core taxa throughout the experiment. The persistence of these characteristics, even during late insular disruption phases, implies that the operation of the LDL has activated an ecological memory mechanism, which progressively homogenized species pools. Although the metacommunity also underwent slow and gradual community succession, its permanent, synchronized structure remained intact and was resistant to invasive bacterial species.

Overall, the results demonstrate that the LDR provides a powerful strategy for the long-term stabilization of highly complex communities. Rather than requiring detailed control of individual species and their interaction, controlled connectivity can maintain multiple local communities in a synchronized and functionally stable state over exceptionally long periods. The LDR provides a framework for long-term management of complex microbial communities in biotechnological settings and for other microbiome-based applications.

## Acknowledgements

This work was supported by Chinese Scholarship Council (CSC) and the German Research Foundation (DFG) within the priority program SPP 2389 ‘Emergent Functions of Bacterial Multicellularity’ Grants SM (PI1) 503905203 and GA (PI2) 503905203.

## Author ORCIDS

Gorkhmaz Abbaszade http://orcid.org/0000-0001-8451-9579 Kathrin Stückrath http://orcid.org/0000-0002-4862-175X Susann Müller http://orcid.org/0000-0003-2060-6256

## Author contribution

Ruyu Gao: Experimentation, Data analysis, Data curation, Data validation, Data visualization, Editing I Gorkhmaz Abbaszade: Data analysis, Data curation, Data validation, Data visualization, Writing, Review, and Editing I Shuang Li: Conceptualization, Experimentation, Data analysis, Data curation, Data validation, Editing I Florian Schattenberg: Experimentation, Data analysis, Review and Editing I Christine Süring: Experimentation I Kathrin Stückrath: Experimentation, Data analysis I Erik Schönfelder: Experimentation, Data analysis I Volker Grimm: Conceptualization, Review and Editing, Susann Müller: Conceptualization, Supervision, Data analysis, Data validation, Writing, Review and Editing.

## Data availability

The flow cytometric data and all RScripts are available on Zenodo: https://zenodo.org/records/24311916. Raw metagenomic sequencing reads have been deposited in the NCBI SRA under BioProject accession PRJNA1494921.

## Additional files

The following material is available online.

Supplementary Information: Section 1 to Section 12

